# Comparing optimized exoskeleton assistance of the hip, knee, and ankle in single and multi-joint configurations

**DOI:** 10.1101/2021.02.19.431882

**Authors:** Patrick W. Franks, Gwendolyn M. Bryan, Russell M. Martin, Ricardo Reyes, Steven H. Collins

## Abstract

Exoskeletons that assist the hip, knee, and ankle joints have begun to improve human mobility, particularly by reducing the metabolic cost of walking. However, direct comparisons of optimal assistance of these joints, or their combinations, have not yet been possible. Assisting multiple joints may be more beneficial than the sum of individual effects, because muscles often span multiple joints, or less effective, because single-joint assistance can indirectly aid other joints. In this study, we used a hip-knee-ankle exoskeleton emulator paired with human-in-the-loop optimization to find single-joint, two-joint, and whole-leg assistance that maximally reduced the metabolic cost of walking for three participants. Hip-only and ankle-only assistance reduced the metabolic cost of walking by 26% and 30% relative to walking in the device unassisted, confirming that both joints are good targets for assistance. Knee-only assistance reduced the metabolic cost of walking by 13%, demonstrating that effective knee assistance is possible. Two-joint assistance reduced the metabolic cost of walking by between 34% and 42%, with the largest improvements coming from hip-ankle assistance. Assisting all three joints reduced the metabolic cost of walking by 50%, showing that at least half of the metabolic energy expended during walking can be saved through exoskeleton assistance. Changes in kinematics and muscle activity indicate that single-joint assistance indirectly assisted muscles at other joints, such that the improvement from whole-leg assistance was smaller than the sum of its single-joint parts. Exoskeletons can assist the entire limb for maximum effect, but a single well-chosen joint can be more efficient when considering additional factors such as weight and cost.

## Introduction

Lower limb exoskeletons can assist human locomotion by reducing the metabolic cost of walking. These devices have the potential to restore ambulatory ability lost from age or disability, and increase maximum performance for high-activity users like first responders, military personnel, or athletes. One important way exoskeletons can help is by reducing the metabolic cost of walking, which is the amount of biochemical energy consumed to produce walking at a given speed (1). Humans tend to move in ways that minimize metabolic cost (2–5), indicating its importance. Reducing the metabolic cost of walking is considered the gold standard for evaluating exoskeletons (6, 7). By reducing metabolic cost, exoskeletons could help achieve related mobility outcomes like increasing the user’s walking speed or decreasing fatigue.

Exoskeletons have improved walking by reducing metabolic cost, but larger improvements may be necessary for widespread adoption of exoskeleton products. Reductions in metabolic cost have been demonstrated from assisting one or two joints (7–19), with the largest metabolic cost reduction being around 18% relative to walking in no exoskeleton (14, 17) and 24% relative to walking in an exoskeleton with no torques applied (12). Despite demonstrated improvements, exoskeletons are in a nascent stage of commercial development, and widespread adoption has not yet occurred (6). To promote adoption, devices may need larger improvements that offset the negative impacts of exoskeletons, such as worn mass, bulkiness, or cost. Wearing a device imposes a metabolic penalty from added mass which could eliminate small benefits. Users may not be able to sense if the exoskeleton is assisting them because the largest improvements in the field are similar to the just-noticeable difference for metabolic cost (around 20%) (20). Improving our scientific and engineering understanding of exoskeleton assistance could deliver larger benefits that may lead to widespread adoption of these devices.

Whole-leg exoskeleton assistance could produce the largest improvements to walking performance. Assisting all of the lower-limb joints simultaneously seems likely to yield the largest energy savings because the hips, knees and ankles all significantly contribute to biological energy consumption during walking (21, 22). Simulations of exoskeleton assistance also indicate that whole-leg devices should be most effective (23–25). Unfortunately, there has been limited testing of whole-leg assistance for able-bodied users (26), and these devices have not yet reduced metabolic cost (27). With improvements to exoskeleton hardware and control, larger benefits might be realizable.

While whole-body assistance may produce larger benefits, single-joint assistance may be more efficient. Assisting just one joint could lead to a smaller, lighter and more cost-effective device, which could result in better net improvements. While a variety of single-joint devices have been tested, there has not been a well-controlled comparison between them owing to differences in actuator capabilities and control. Each device has had different limits on torque and power (28), many below previously-identified optimal values (12), and larger torque and power capacity are associated with larger reductions in metabolic cost (13, 14). Most devices have not used control that has been systematically optimized for the participant, which can improve performance by as much as a factor of five (12, 29). A direct comparison of optimized single-joint assistance in a high-torque, high-power exoskeleton would be useful to designers as they choose which joint, and how many joints, to assist.

A comparison of single-joint and multi-joint assistance would also provide scientific insights into the biomechanics of walking. A large portion of human leg musculature is biarticular (30), comprising muscles that span two joints. Biarticular muscles might be more effectively assisted by multi-joint exoskeletons, leading to a total benefit beyond the sum of assisting each joint individually. Some experiments have suggested that multi-joint assistance might be more effective than single-joint assistance (16, 31). Alternatively, adding assistance at other joints may have diminishing returns. Users might adapt their walking pattern to maximize the benefit from assistance at a single-joint, thereby indirectly benefiting muscles at other joints, which may make it relatively less effective to assist additional joints. Some experiments have shown indications of such indirect assistance (32, 33). A well-controlled comparison allowing observations of how users respond to different types of assistance would help us to develop improved models of biomechanical and neural adaptation to exoskeletons.

We recently developed whole-leg exoskeleton hardware and optimization techniques that enable comparisons of assistance at different combinations of lower-limb joints. We built a tethered hip-knee-ankle exoskeleton emulator that can assist hip flexion and extension, knee flexion and extension, and ankle plantarflexion of both legs (28). This device can apply large torques using offboard motors and power, enabling laboratory tests of different assistance strategies without actuation limits (34). We can pair this tool with human-in-the-loop optimization, a process where the control of the exoskeleton is updated in real time based on biomechanical measurements of the user (12, 35–37). This approach has led to the largest improvements in the metabolic cost of walking when paired with exoskeleton assistance (12, 14). By optimizing assistance at one, two, and three joints, we will be able to compare the best possible assistance outcomes for each joint.

The purpose of this study was to identify the maximum possible improvements to the metabolic cost of walking with lower limb exoskeleton assistance and to understand how assistance at each joint contributes to this improvement. We performed experiments optimizing and comparing assistance at the hips, knees, and ankles, individually, in pairs, and with all three joints assisted simultaneously. We optimized each assistance pattern to reduce the measured metabolic cost of walking and compared it to walking without assistance and walking without the exoskeleton. We measured changes in kinematics and muscle activity to see how users adapted to assistance and to gain insights into the potential biomechanical mechanisms that brought about reductions in metabolic cost. By finding and comparing optimized assistance for different potential device architectures, we expected these results to inform models of human adaptation to exoskeletons and lead to the design of more effective exoskeletons.

## Results

### Human-in-the-loop optimization

We performed three different experiments optimizing exoskeleton assistance from the hip-knee-ankle exoskeleton (Fig. 1). We first optimized hip-only, knee-only, and ankle-only assistance to compare the best possible assistance of each single joint (N = 3, 1F 2M, 61 - 90 kg, age 19-26). We next optimized two-joint assistance in different combinations to evaluate the benefit of adding a second joint (N = 1, M, 90 kg, 26). Finally, we optimized whole-leg assistance of the hip, knee and ankle to try to identify the maximum reduction in metabolic rate that exoskeleton assistance can provide during walking (N = 3, 1F 2M, 61 - 90 kg, age 19-26, same as single-joint experiment). For all conditions, the algorithm minimized the measured metabolic cost of walking at 1.25 m/s. Two participants had previous experience in the exoskeleton, and the third completed a lengthy training protocol before optimization (Sup. Note 1).

**Fig 1.**
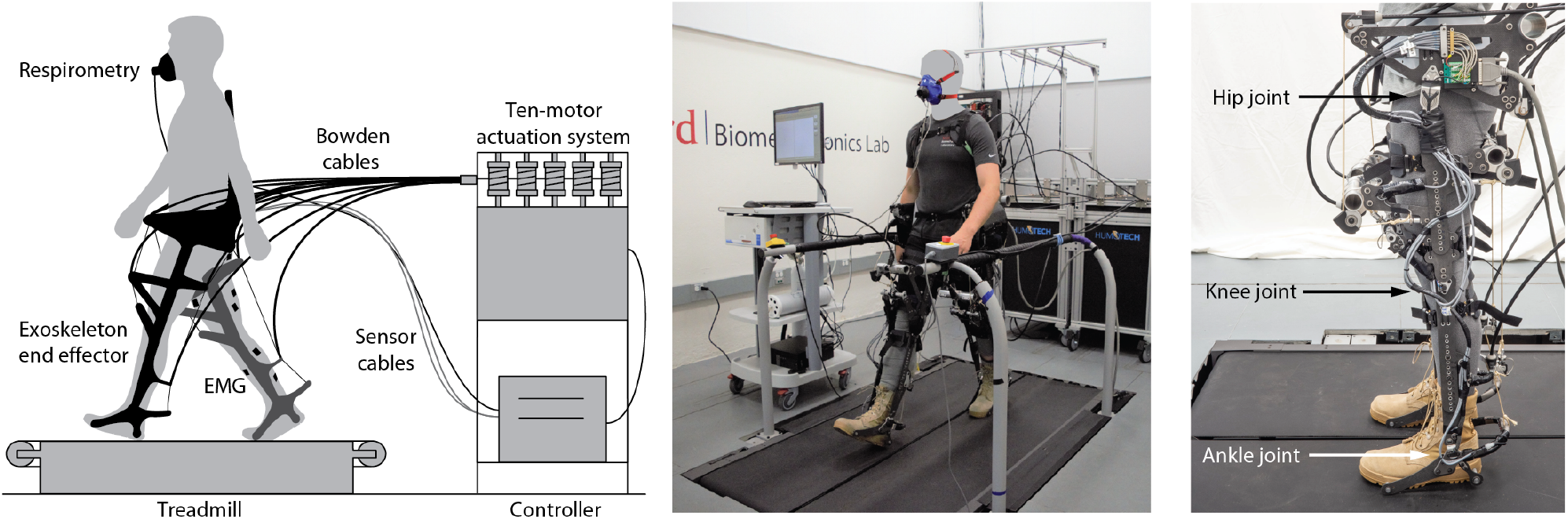
Overview of exoskeleton emulator system. (Left) Overview of exoskeleton emulator. Ten powerful off-board motors actuate a lightweight end effector worn by a user who walks on a treadmill. Metabolic cost is measured using a respirometry system and muscle activity is measured using electromyography (EMG). (Center) Isometric photo of experimental setup. (Right) Side view of exoskeleton. The exoskeleton can apply torques in hip flexion and extension, knee flexion and extension, and ankle plantarflexion.

### Metabolic cost

The metabolic cost of walking was reduced in all assistance conditions (Fig. 2). To evaluate the metabolic cost of walking, we subtracted the metabolic cost of standing quietly (1.55 W/kg on average, Sup. Note 2) from the metabolic cost measured in each walking trial. The metabolic cost of walking without wearing the exoskeleton was 2.91 W/kg on average. The cost of walking in the exoskeleton with no torque applied was 3.70 W/kg on average, which was higher than the no-exoskeleton condition because of the added mass and impedance to the user. Percent reductions in metabolic cost were calculated in comparison to this no-torque condition to assess the effect of the designed torque assistance specifically, allowing for best comparisons between assistance conditions.

**Fig 2.**
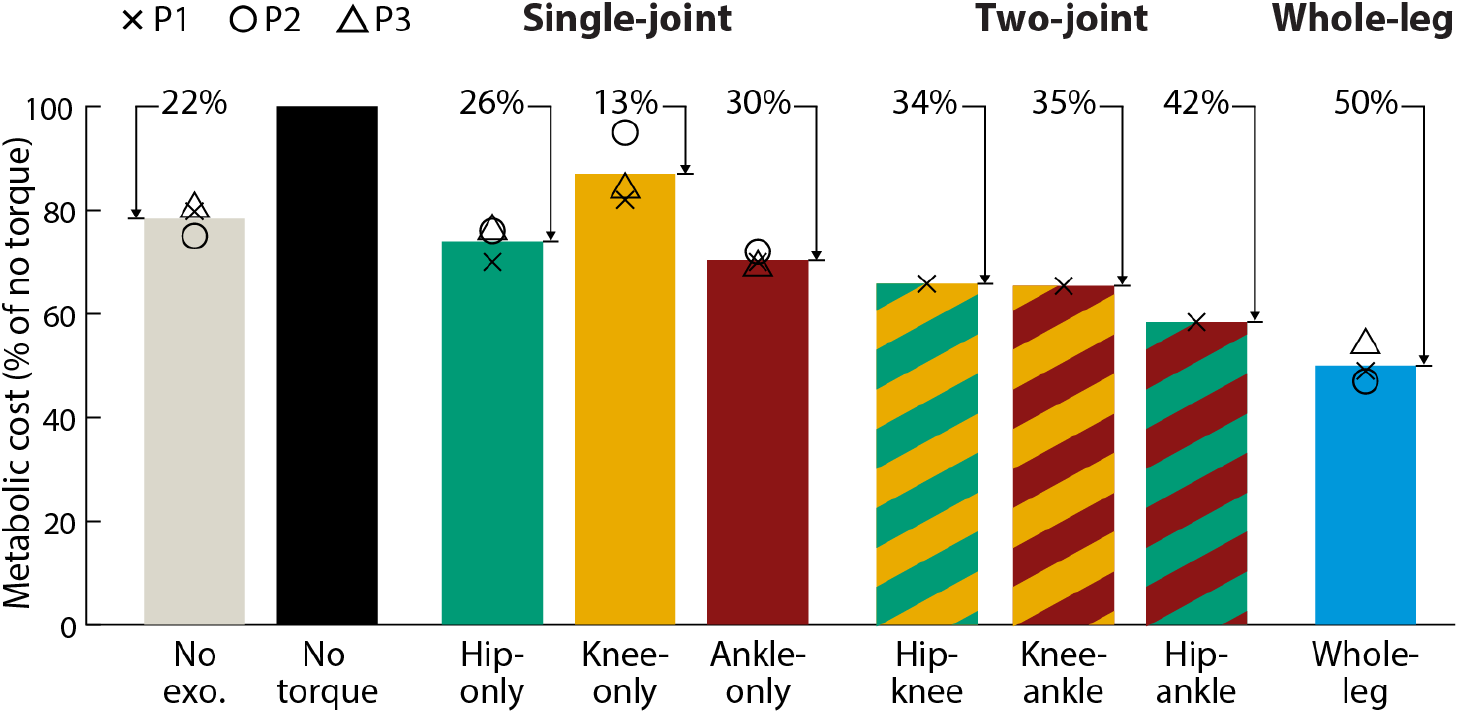
Metabolic cost of walking. Average metabolic cost (bar) of each condition reported as a percentage of walking in the exoskeleton with no torque. Individual participant values are shown with symbols (p1 X, p2 O, p3 Δ). Metabolic cost of walking was calculated by subtracting out quiet standing. The percent reduction relative to walking with no torque is shown above each bar. For each participant, the cost of walking without the exoskeleton (No exo., gray) was averaged over all validations. Two-joint assistance (striped, two-colors) was optimized for one participant. Whole-leg assistance (blue) provided the largest improvement to metabolic cost of walking, reducing it by 50% relative to walking in the exoskeleton without assistance.

Single-joint assistance at each joint reduced metabolic cost, with the largest improvements coming from the ankles and hips. Hip-only assistance reduced the metabolic cost of walking by 26% relative to walking with no torque (N = 3, range of reductions: 24% - 30%, p = 0.005). Knee-only assistance reduced metabolic cost for each participant, with an average reduction of 13% relative to walking with no torque, although this was not statistically significant (N = 3, range of reductions: 5% - 18%, p = 0.07). Ankle-only assistance performed best of the single-joint strategies, reducing metabolic cost by 30% relative to walking with no torque (N = 3, range of reductions: 28% - 31%, p = 0.004). When assisting a single joint, exoskeleton designers should consider the ankles or the hips.

Two-joint assistance outperformed single-joint assistance. Two-joint assistance reduced metabolic cost of walking relative to no torque by 42%, 35%, and 34% for hip-ankle, knee-ankle and hip-knee assistance, respectively (N = 1, Sup. Note 3). Hip-ankle assistance provided the most benefit, mirroring single-joint reductions and aligning with expectations based on biological power.

Whole-leg exoskeleton assistance led to the greatest reductions in metabolic cost of any condition. Whole-leg assistance (hips, knees, and ankles simultaneously) reduced the metabolic cost of walking by 50% relative to walking with no torque (N = 3, range of reductions: 46% - 53%, p = 0.02), corresponding to a reduction of 37% relative to walking without wearing the exoskeleton (N = 3, range of reductions: 34% - 41%, p = 0.02). This shows that about half of the metabolic energy expended during walking can be saved through exoskeleton assistance, and suggests that whole-leg assistance could provide large net benefits in untethered systems, even after accounting for the effects of added mass.

### Optimized exoskeleton torque

Optimized torques differed from biological torques in both timing and magnitude (Fig. 3, Sup. Note 4, Sup. Note 5, Sup. Note 6). Optimized torque magnitudes were smaller than biological torques, with peak exoskeleton torques ranging from about 15% to 60% of biological peaks. Ankle torque magnitudes were largest and optimized to the comfort-limited parameter constraints in all but one case. The timing of the optimized assistance only partially aligned with biological torque. For example, the peak of optimal hip flexion assistance occurred later than peak biological flexion torque. Sometimes, assistance torque opposed typical biological torques. For example, knee flexion assistance around 60% of stride opposed biological knee extension torque for normal walking. These optimized magnitudes indicate the design requirements for mobile devices and show that optimized assistance is not a scaled version of biological torques.

**Fig 3.**
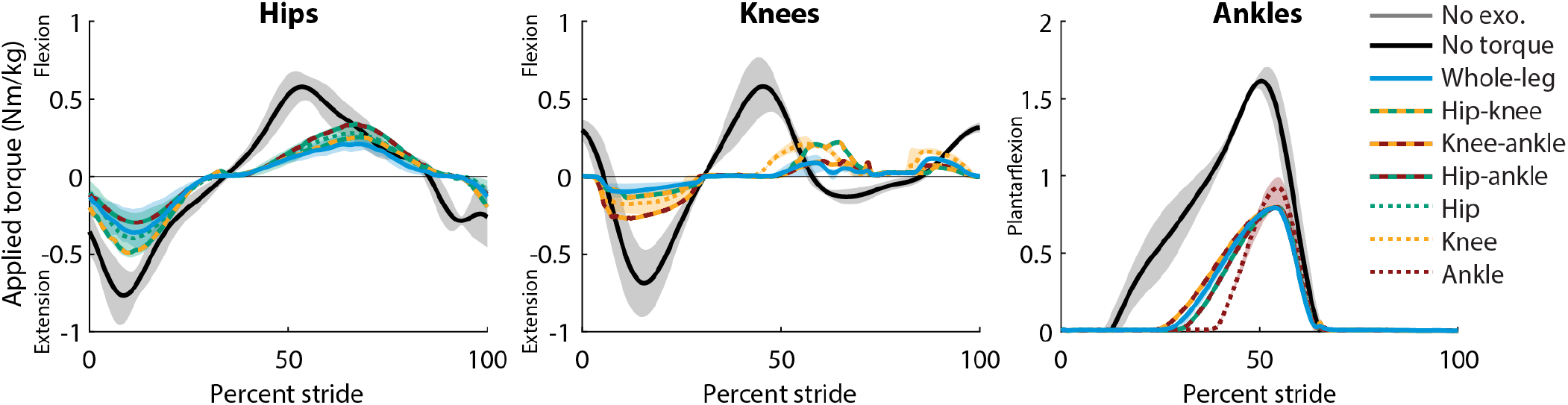
Optimized exoskeleton torques. Optimized single-joint (single-color, dotted), two-joint (two-color, dashed) and whole-leg (blue, solid) exoskeleton assistance torques at the hips (left), knees (center), and ankles (right). Lines are the average of the measured applied torque profiles across both legs and all participants (N = 3 for single-joint and whole-leg, N = 1 for two-joint), with the range of optimized profiles shown with their respective clouds for single-joint and whole-leg assistance. Biological joint torques for unassisted walking without an exoskeleton (black) are included from a different study with different participants (25, 38) for reference; gray clouds indicate standard deviation of biological torques. For the hips and knees, whole-leg assistance optimized to smaller magnitudes than single-joint assistance. For the ankles, maximum torque had to be constrained to find comfortable profiles for walking. Ankle torques were limited to 1 Nm/kg for single-joint assistance, and 0.8 Nm/kg for two-joint and whole-leg assistance.

The shape and timing of optimized assistance was consistent across conditions and participants, but optimal magnitudes differed. For example, the optimal timing of peak hip extension assistance was about 11% of stride for all joint combinations and participants, while optimal magnitudes ranged from Nm/kg (hip-ankle) to 0.5 Nm/kg (hip-knee). One exception to the consistency of optimal timing was ankle torque rise time, which was shorter during single-joint assistance, possibly due to adjustments for comfort in multi-joint conditions (Sup. Note 5). Optimized assistance torques were typically larger when acting alone at a joint, and smaller when acting in a multi-joint configuration. For example, for P1, applied knee flexion torque peaked at 0.25 Nm/kg for knee-only assistance and at 0.14 Nm/kg for whole-leg assistance. The consistency of optimal timing parameters suggests that optimization could occur in a lower-dimensional parameter space of torque magnitudes, and that a generalized assistance profile could be almost as effective as a customized one.

### Kinematics

Kinematics varied between assistance conditions, indicating that the user’s walking pattern is not fixed and adapts to best utilize assistance (Fig. 4, Sup. Note 7, Sup. Note 8). These changes were beyond the deviation measured during walking with no torque (gray cloud, Fig. 4). In some cases, assistance shifted joint angles in the direction of the applied torque. For example, peak ankle plantarflexion angle increased with whole-leg assistance, and increased even more during ankle-only assistance, which had larger ankle torques. However, some kinematic changes were not the direct result of applied torques. For example, the indirect effects of hip-only and ankle-only assistance on the knee during stance were larger than the direct effect of knee assistance. These kinematic adaptations indicate the user adjusts their walking strategy to maximize the benefit they get from the exoskeleton, and that these adaptations don’t always match intuition.

**Fig 4.**
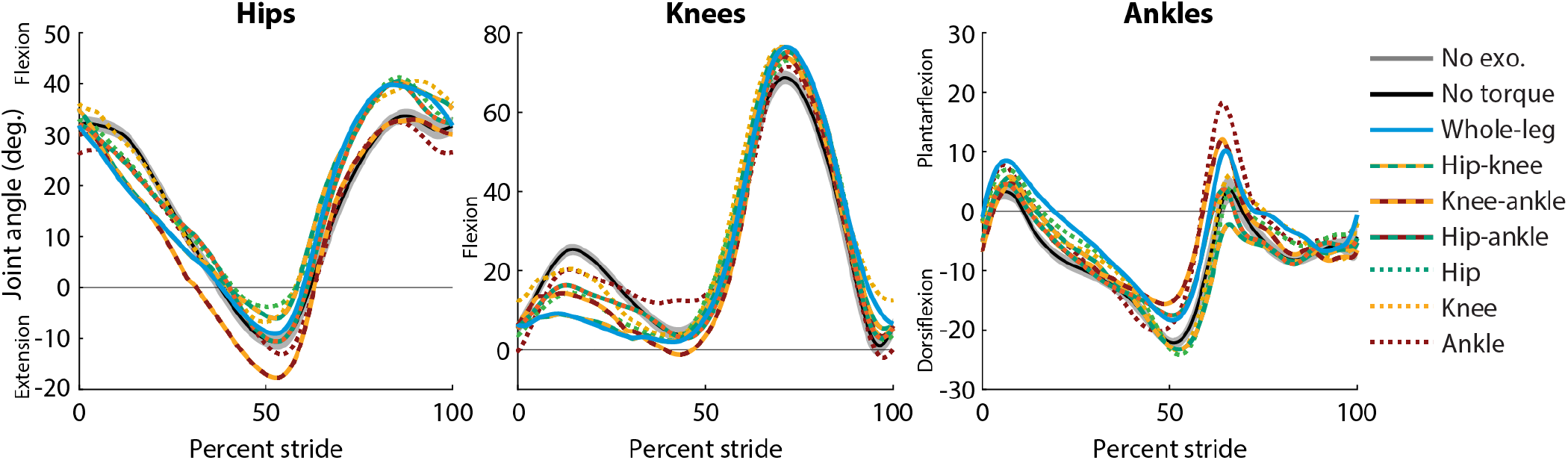
Average joint kinematics. Average joint angle as a percentage of stride at the hips (left), knees (center), and ankles (right) for each condition. Shown here are the average for both legs across all participants (N = 3 for single-joint and whole-leg, N = 1 for two-joint). All single-joint and whole-leg conditions were tested on the same day to reduce changes in alignment between user and device. Two-joint conditions were each collected individually. For walking in the exoskeleton with no torque (black), the standard deviation of angles is shown (gray cloud) to contextualize the magnitude of changes between conditions.

### Muscle activity

Muscle activity decreased with assistance, but it was not completely eliminated (Fig. 5, Sup. Note 9). Typically, reductions in activity were seen in muscles that crossed assisted joints and acted in the same direction as assistance. For example, soleus activity decreased during all conditions that applied ankle assistance. Sometimes activity increased during opposing assistance, such as in the vastus lateralis during periods of knee flexion torque. Some reductions in muscle activity occurred during assistance at other joints. For example, gluteus maximus activity decreased when the hip was assisted directly, but also decreased during ankle-only assistance. This is consistent with the observation that exoskeleton assistance at one joint can indirectly assist muscles that cross other joints. The indirect assistance could be from the complex dynamics of the leg during walking, or could be facilitated by the kinematic adaptations to maximize the effectiveness of each type of assistance. Optimized assistance did not cause users to eliminate muscle activity, suggesting that either some amount of activity is still useful or further advancements to control architecture would be needed to reduce energy expenditure further.

**Fig 5.**
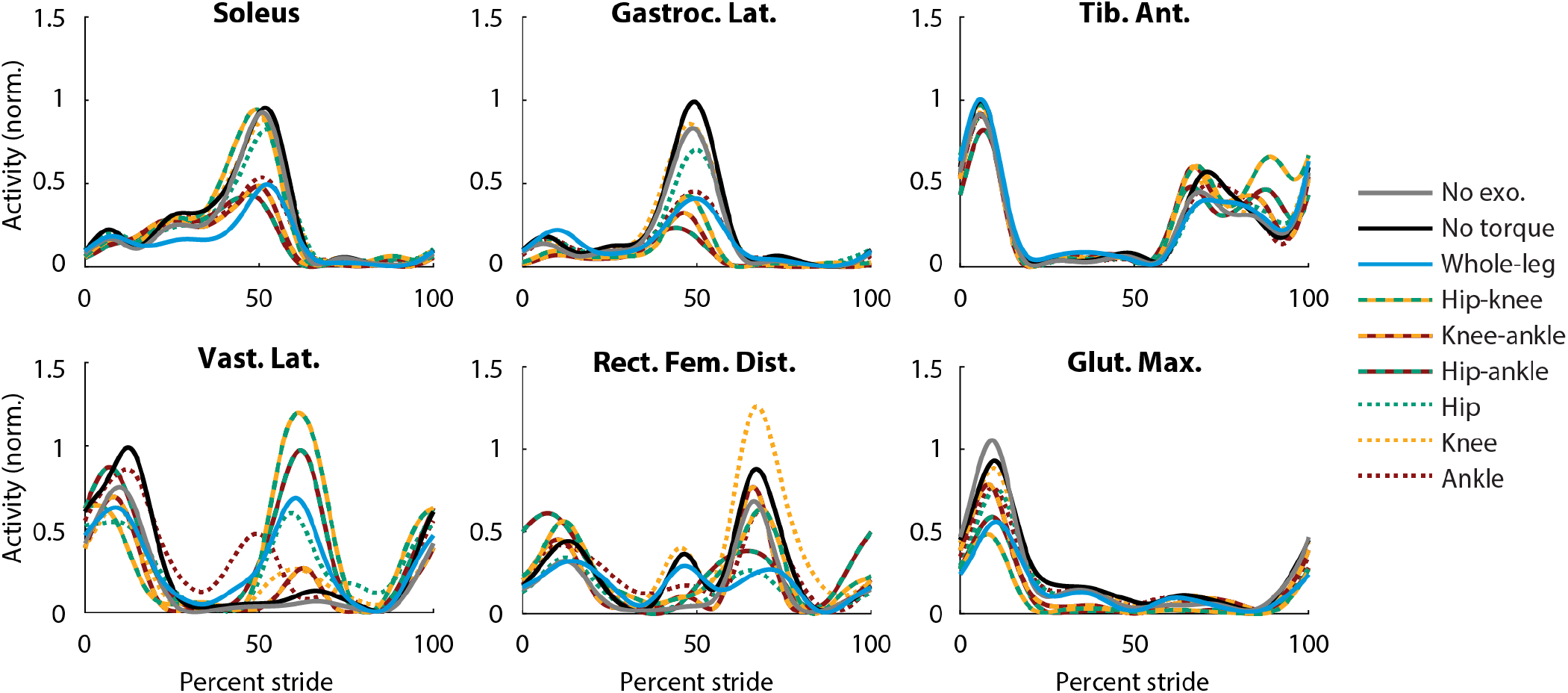
Muscle activity. Muscle activity measured during walking using surface EMG for each condition. Lines shown are the average across all participants (N = 3 for single-joint and whole-leg, N = 1 for two-joint). The EMG signal was filtered, averaged, had baseline activity removed to eliminate noise, and normalized to the peak value of walking in the exoskeleton without assistance (black). Gluteus maximus activity (second row, third column) decreased for hip-only, hip-knee, hip-ankle, and whole-leg assistance as expected, and also decreased during ankle-only assistance, indicating that the gluteus was indirectly assisted by ankle exoskeleton torque. This effect was less pronounced for the soleus (top row, first column), where hip-only assistance only slightly reduced muscle activity.

## Discussion

With capable devices, optimization, and training, exoskeletons can provide very large improvements in locomotor performance. Whole-leg assistance reduced the metabolic cost of walking by 50% relative to walking with no torque, a sub-stantial improvement over the state-of-the-art (17% - 24%) (12, 14, 17)). This corresponded to a 37% reduction relative to walking with no exoskeleton, nearly double the just-noticeable difference in metabolic cost (20%) (20), indicating that participants could feel the reduction in effort compared to walking normally. Because whole-leg assistance produced the largest benefit of all assistance conditions, exoskeleton designers who want to maximize performance should consider assisting the whole leg.

Among single-joint assistance strategies, ankle-only assistance was most effective, followed closely by the hip, with smaller reductions possible at the knee. Hip and knee assistance resulted in greater metabolic cost reductions than previous lower-torque exoskeletons assisting these joints (14, 17, 18), while ankle assistance resulted in similar improvements as found with high-torque exoskeletons and human-in-the-loop optimization (12, 39). Knee-only assistance reduced the metabolic cost of walking, although the reduction was not statistically significant for our sample size and significance level. Knee-only assistance may be more effective for walking up inclines (18, 40), considering the increased positive power requirements from the knee (41). Devices designed to assist just one joint during level walking should target the ankle or hip, which showed reductions of 30% and 26%, respectively. With similar metabolic reductions, designers could compare between the ankle or hip based on other aspects of the design. For example, it may be easier to interface with the ankle using a simple device, while a hip exoskeleton places the mass more proximally on the body where it is easier to carry (42).

The best-performing two-joint assistance strategy, hip-ankle assistance, reduced metabolic cost by 42%, nearly as much as whole-leg assistance. With 35% and 34% reductions for knee-ankle and hip-knee assistance, the small added benefit of knee-assistance may not be worth the added device complexity compared to ankle-only or hip-only assistance for level ground walking. The inclusion of knee assistance may be more effective for different walking conditions such as incline walking or during sit-to-stand, where the knee is expected to be contributing more to movement.

Assisting multiple joints results in larger net benefits, but smaller benefits per joint, possibly because of the way people adapt to exoskeleton assistance. Whole-leg assistance led to the largest metabolic cost reduction (50%), but it was smaller than the sum of the reductions of the single-joint assistance strategies (26% + 13% + 30% = 69%). It appears that the benefits of indirectly assisting muscles at other joints outweigh those of directly assisting bi-articular musculature. The reduction in metabolic cost per joint assisted (Table 1) could be helpful for designers when considering potential device architectures.

**Table 1.** Metabolic reduction per joint assisted, relative to walking in the exoskeleton with no torque.

| Condition | Ankle-only | Hip-only | Hip-ankle | Knee-ankle | Hip-knee | Whole-leg | Knee-only |
| --- | --- | --- | --- | --- | --- | --- | --- |
| Reduction per joint | 30% | 26% | 21% | 17% | 17% | 17% | 13% |
| Total reduction | 30% | 26% | 42% | 35% | 34% | 50% | 13% |

These results suggest ways of designing better exoskeleton products. Optimized torques did not mimic biological torques, with magnitudes smaller than biological for all joints and peak torques later than biological peaks for the hips and ankles (Fig. 3). Optimized torque magnitudes were within the range of reported capabilities of some existing mobile devices for the lightest participant (61 kg) (9, 15, 43) but not for the heaviest participant (90 kg) (28). Exoskeletons could be designed in different sizes (e.g. small, medium, large) that meet the optimized torque magnitudes for different sized users while minimizing worn mass. The similar timing parameters across users and across assistance strategies suggest that these optimized profiles could translate well to existing devices with lower capabilities and could be generalizable to a wide range of users. Device designers could consider control strategies that allow for kinematic adaptations because they seem to be useful to maximize device effectiveness. We also recommend considering state-based control at the knees, which was more effective than pilot tests of strictly torque-time control. This strategy may have facilitated adaptation to knee extension assistance during stance, because the torques grew as the user bent more into flexion, allowing for a “stabilizing” effect that prevented buckling of the knee. These findings should translate well to a whole-leg mobile device because the worn mass (13.5 kg) is similar to the expected mass of a mobile device capable of the optimized torque magnitudes (10 kg) based on published torque densities (9, 11, 28, 43–48) (Sup. Note 10).

These results can be used to improve our models of human coordination, especially when using assistive devices. The larger metabolic cost reductions we saw from hip and ankle assistance support the idea that the hips and ankles are primary energy consumers during walking (22). Unlike some simulations (25), muscle activity did not go to zero even when assisted without hitting torque limits, indicating the user is optimizing for more than just metabolic cost for an average steady-state stride. Simulations could capture that more complicated objective function, including control required for balance. These results show that kinematic adaptations to assistance are important and should be considered in simulations.

This study could have been improved by testing more participants, providing additional training, or testing additional controller parameterizations. This was an extremely arduous experiment with long optimization times, with each participant completed over 50 hours of experiments. It was then interrupted by difficult external conditions (the COVID-19 pandemic). As such, we were only able to complete three participants for the single-joint and whole-leg optimizations, and one participant for the two-joint optimizations. However, given the magnitude of the changes and the consistency of the responses across participants, this sample size is sufficient to identify the efficacy of the joint combinations tested. Given three participants and a desired statistical power of 0.8, and assuming metabolic reductions have a standard deviation of 7.3% (12), we can confidently detect metabolic reductions of 24% and larger. Although we gave our users substantial training and optimization time, more time may have improved the outcomes. Longitudinal studies with mobile devices that can be worn daily could show greater improvements to walking as users adapt. Torques optimized to the comfort-based limits at the ankles, which were set due to discomfort at the biological ankle joint, possibly from extending the ankles too quickly or too far during torque application at push-off. Using a more sophisticated control approach to ensure user comfort while allowing the largest possible exoskeleton torques might also lead to larger benefits.

These results suggest that new cost functions, gait environments, and user populations could be exciting topics for future studies. Future work could optimize metabolic cost alongside other costs that are important for gait, such as walking speed, balance, or user satisfaction (49). Our study did not penalize high torques or powers, but future work could try to maintain sufficient metabolic cost reductions while minimizing actuator requirements, which can be costly to mobile devices. While our study assisted walking at a fixed speed on level-ground, future work can explore optimized assistance for walking in different conditions such as at different speeds, on inclines, or with worn loads. Our study was restricted to a treadmill due to our tethered device, but this work could be extended to unstructured environments by translating the paradigm to mobile devices. Our findings for assisting young, able-bodied users could be a starting point to optimize assistance for older adults and people with disabilities, hopefully speeding the discovery of effective assistance strategies.

## Methods and Materials

### IRB Approval

All user experiments were approved by the Stanford University Institutional Review Board and the US Army Medical Research and Materiel Command (USAM-RMC) Office of Research Protections. All participants provided written informed consent before their participation as required by the approved protocol.

### Participants

Three healthy participants were included in this study (P1: M, 26 years old, 90 kg, 187 cm; P2: F, 26 years old, 61 kg, 170 cm; P3: M, 19 years old, 82 kg, 176 cm). We were limited to three participants because of the extensive time required to complete the protocol (Sup. Note 11, Sup. Note 12). Each participant completed at least 50 hours of experiments in total. These participants were also authors of the study (PF, GB, RR) as these were the people who could spend such time as a participant for the study. With three participants, we have a statistical power of 0.8 to detect metabolic reductions greater than 24%, assuming metabolic reductions have a standard deviation of 7.4% (12) (Sup. Note 12). With three participants, the 50% reduction detected from whole-leg assistance has a statistical power of 0.999.

All three participants were experienced with the device at the time of optimization. P1 and P2 had previous experience walking in the exoskeleton before this experiment. P3 completed a training protocol prior to optimization to get accustomed to wearing the exoskeleton and walking with torques. More details on the training protocol are included in Sup. Note 1.

### Experimental protocol

We optimized single-joint, two-joint, and whole-leg assistance using a hip-knee-ankle exoskeleton (Fig. 1) (28). We used human-in-the-loop optimization (12), a strategy where the control of the exoskeleton is updated in real-time based on measurements of the user. For this study, the cost function to be minimized was the measured metabolic cost of walking at 1.25 m/s. First, we optimized single-joint assistance for all three users in the order of ankle-only, hip-only, and then knee-only assistance. We then optimized whole-leg assistance for all three users, meaning we optimized assistance of the hip, knee and ankle simultaneously. Finally, for one user (P1), we optimized two-joint assistance, in the order of hip-ankle, knee-ankle, and hip-knee assistance.

After optimization, we performed validation experiments to compare the optimized assistance to the control conditions of walking without the exoskeleton, and walking in the exoskeleton with no torque applied. During these validations we measured metabolic cost, applied torques, kinematics, and muscle activity.

### Exoskeleton hardware

Assistance was applied using a hip-knee-ankle exoskeleton emulator (Fig. 1) (28). The exoskeleton emulator uses powerful off-board motors and Bowden cable transmissions to actuate an end effector worn by the user. The device has a worn mass of 13.5 kg. It has carbon fiber struts along the length of the legs that are designed to minimize restriction of the user by being stiff in actuated directions but compliant in out-of-plane bending. The exoskeleton was fit to each user by adding boots for their foot size, by adjusting the length of the shank, thigh, and torso segments of the exoskeleton, and by adjusting the width of the exoskeleton at the knees, thighs, and hips. Straps were adjusted to fit the user at the shanks, thighs, hips, and torso.

### Exoskeleton control

The exoskeleton is controlled by commanding a desired torque for each joint (28). When the desired torque is zero, the exoskeleton tracks the user’s joint angles and applies no torques. During walking, we define these desired torque profiles as a function of percent stride. We consider heel strike, measured by ground reaction forces in the treadmill, to be the start of a stride. We calculate percent stride as the time since heel strike divided by the average stride time over the past 20 strides. The hip profile starts 84% of stride after heel strike because hip extension torque is active during heel strike. Resetting the hips’ stride time at heel strike caused discrete jumps in hip extension torque during pilot testing. The desired torque profile for each joint is made up of a spline (piecewise cubic hermite interpolating polynomial) anchored by nodes. Each node can be set in advance by an operator, or it can be updated in real time by an algorithm. For the knees, torque was also commanded as a function of joint state. Along with a torque-time profile, knee torque had one spring-like phase during stance, and one damping-like phase during late swing.

Our exoskeleton accurately applied desired torques using closed-loop proportional control with iterative learning and joint-velocity compensation (28, 50). Root-mean-square (RMS) error for tracking desired torques was 0.6 Nm at the hips, 3.0 Nm at the knees, and 0.4 Nm at the ankles during whole-leg assistance (Sup. Note 13). Error was highest at the knees because the state-based control allowed for discontinuous jumps in desired torque that were not possible for our device to track, and the desired torque would change step to step making it harder to track. When zero torque was commanded it was realized effectively, with an RMS applied torque of less than 1 Nm.

### Controller parameterization

The optimization algorithm varied parameters that affected the desired torque control of the exoskeleton (Fig. 6). These parameters are mostly related to the timing and torque magnitude of the nodes that define our splines.

**Fig 6.**
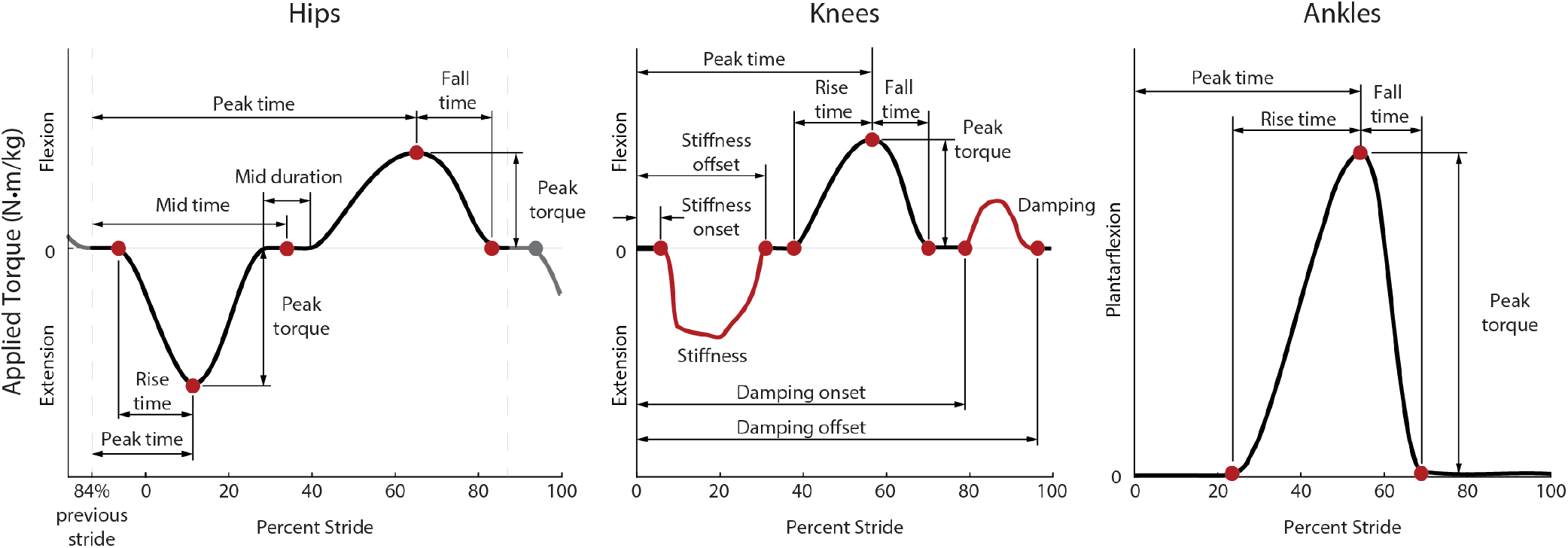
Parameterization of applied torque. For the hips (left) and ankles (right), torque (black) was commanded as a function of time, defined as a spline fit to nodes (red) that were optimized during the experiment. For the knees (center), torque was commanded both as a function of time (black), joint angle, and joint velocity. During stance, the knee torque was a function of knee angle to mimic a spring (red), where the spring’s stiffness was optimized. During late swing, torque was a function of knee joint velocity to mimic a damper (red). Hip-only assistance was determined by 8 parameters, knee-only assistance had 10 parameters, ankle-only assistance had 4 parameters, and whole-leg assistance optimized all 22 parameters.

We chose these parameters by considering previously successful human-in-the-loop optimizations (12, 14), considering biological torques during walking (21), and by pilot testing. Before this, one participant completed a 9-parameter whole-leg optimization pilot study (Sup. Note 14), which indicated the need for more degrees of freedom in our controller.

For these optimizations, we included 8 parameters for the hips, 10 for the knees, and 4 for the ankles (Fig. 6). For optimization of whole-leg assistance, the optimizer could adjust all 22 parameters. Each parameter had a minimum and maximum allowed value. The allowed parameter ranges were based on user testing to ensure all tested profiles would be sufficiently comfortable for the user to walk in. The optimization tended not to optimize to these limits. However, ankle torque was often as large and as late in stride as possible, which meant the fall time was minimized to prevent torque application during swing. Tables with the parameter ranges, as well as their initial and optimized values, are available in Sup. Note 5.

The hip profile was defined by 8 parameters. Hip extension was defined by the rise time, peak time and peak magnitude while hip flexion was defined by the peak time, peak magnitude and fall time. There was a period of no torque in between the two peaks defined by the midpoint timing and its duration. The period of no torque dictated the hip extension fall time and the hip flexion rise time.

Knee torque was defined by 10 parameters. Knee torque was commanded both as a function of percent stride and of joint state. The first phase of knee torque was knee extension defined as a virtual spring, with torque proportional to knee angle, which was zero when the knee was straight. Knee extension was defined by the virtual spring onset timing, stiffness, and offset timing. If the joint angle reached zero degrees before the offset time, the knee torque would stay at zero torque for the remainder of the stiffness period. During knee flexion around toe-off, torque was defined as a function of time similar to the hips. This torque was determined by the rise time, peak torque magnitude, peak time, and fall time. Late in swing, knee flexion torque was commanded as a virtual damper, so torque was proportional to a filtered measurement of knee joint velocity. The damping period was defined in a similar way to knee stiffness, with optimization of the onset timing, the damping coefficient, and the offset timing. As the knee joint angle and velocity were not necessarily zero at the start of the state-based controllers, desired torque could instantaneously change at the onset.

Ankle torque was defined using 4 parameters which were previously effective for optimization of ankle assistance (12). Torque was defined by rise time, peak torque magnitude, peak time, and fall time. To ensure large torques weren’t applied too late in the stride, torque was set to be zero by 65% of stride, so if peak time optimized to it’s latest allowed value (55% of stride), the fall time would be set to the minimum allowed fall time (10% of stride).

### Human-in-the-loop optimization protocol

To optimize assistance, we used the covariance matrix adaptation evolutionary strategy (CMA-ES) (51), which has previously been effective for human-in-the-loop optimization of exoskeletons (12, 52). CMA-ES samples a “generation” of conditions from a distribution defined by parameter means and a covariance matrix, ranks the performance of the samples, and uses those results to update the mean and covariance before sampling the next generation. The optimizer’s goal was to minimize metabolic cost, which was estimated for each condition after two minutes of walking using a first-order dynamical model (53), similar to previous work (12, 52). More details about the optimization, including hyperparameters and numbers of conditions per generation, are included in Sup. Note 15.

The initial parameter values for each optimization were carefully selected to try to reduce convergence time. For the single-joint optimizations for P1, initial parameter values were based on previously optimized assistance (12, 14), hand-tuning, and a 9-parameter pilot study (Sup. Note 14). For whole-leg optimization for P1, initial values were based on the optimized values for single-joint assistance. For P2 and P3, initial values for all optimizations were based on the optimized values for P1. Finally, for the two-joint assistance optimizations, initial parameters were based on the optimized whole-leg assistance values for P1. The initial values for all parameters are included in Sup. Note 5.

The optimization time was intended to balance being long enough to ensure convergence while short enough to be experimentally feasible. P1 underwent a longer optimization to ensure convergence, to estimate expected reductions, and to inform our understanding of how the optimizer would perform (Sup. Note 11, Sup. Note 16). Ankle-only optimization was conducted for twelve generations over three days, and hip-only, knee-only and whole-leg were conducted for at least nine generations over three days. Each two-joint assistance optimization was conducted for six generations over two days. For P2 and P3, single-joint and whole-leg optimization each occurred over at least three days, which seemed sufficiently long for P1 to reach metabolic reductions that were consistent across days and that matched previous studies for previously-assisted joints (12) (Sup. Note 11). For whole-leg assistance for P3, the optimization was restarted because the user had an abnormally high stride frequency that had high metabolic cost, indicating a maladaptation to assistance similar to some users in a previous optimization study. The exact number of generations and days for each optimization is included in Sup. Note 11. Participants were permitted but not required to take breaks between generations. While walking, participants were allowed to listen to podcasts using wireless headphones.

### Validation Protocol

We conducted validation experiments to evaluate the effectiveness of the optimized assistance. Metabolic reductions were validated for each assistance strategy after each optimization. After all the single-joint and whole-leg optimizations were completed, torques, kinematics, and muscle activity were compared between assistance strategies on the same day. Finally, the two-joint assistance strategies were validated after each optimization.

After each optimization was completed, a validation experiment was used to accurately assess the metabolic cost of walking and calculate the percent reduction. This collection was on a separate day, before optimization of the next assistance strategy began. Users walked in longer bouts for the exoskeleton conditions to ensure accurate measurements of steady-state metabolics and to ensure users were adapted to the device and assistance. We recorded the user standing quietly for 6 minutes, walking without the exoskeleton for 6 minutes, walking in the exoskeleton with no torque applied for 10 minutes, and walking with the optimized assistance torques for 20 minutes, in a double-reversed order (ABCDD-CBA). This order was not randomized due to the time it takes to get in and out of the exoskeleton, as well as to maximize acclamation to the device by presenting progressively more novel conditions. Users rested for at least three minutes between walking conditions, and at least five minutes before a quiet standing condition, to ensure their metabolics returned to baseline. For the no exoskeleton condition, users wore the same brand and model of boots that are included in the exoskeleton (McRae 8189).

After all single-joint and whole-leg optimizations were completed, we evaluated all these optimized strategies in one data collection to directly compare conditions. We measured applied torque, kinematics, muscle activity, power, and vertical ground reaction forces. For this validation, we recorded the user standing quietly for six minutes, walking without the exoskeleton for six minutes, walking in the exoskeleton with no torque applied for ten minutes, and then walking for ten minutes in each of the four optimized assistance conditions (hip-only, knee-only, ankle-only, whole-leg) in a random order, and then the no torque, no exoskeleton, and quiet standing conditions a second time (ABCDEFGCBA).

For the one user who completed optimization of two-joint assistance, a validation day was completed after each optimization similar to the protocol following the single-joint and whole-leg optimizations. We recorded the user standing quietly for six minutes, walking without the exoskeleton for six minutes, walking in the exoskeleton with no torque applied for ten minutes, and walking with the optimized assistance torques for 20 minutes, in a double-reversed order (ABCDD-CBA), so each condition was evaluated twice. For these validations, we measured metabolics, torques, kinematics, muscle activity, power, and vertical ground reaction forces.

### Measured Outcomes

We collected biomechanical data of the user when walking in the different conditions during the validation. We calculated the average of these measurements over the last three to five minutes of walking of each condition to ensure the user’s metabolics and gait had reached steady-state.

#### Metabolic Cost

Metabolic cost was calculated using indirect calorimetry. We measured volumetric carbon dioxide expulsion, oxygen consumption, and breath duration on a breath-by-breath basis (Quark CPET, COSMED). For each condition, we calculated metabolic rate using a modified Brockway equation (54) similar to previous studies (12, 52). Average metabolic cost was calculated for each condition using the last three minutes for quiet standing and walking with no exoskeleton, and using the last five minutes for walking with no torque and walking with optimized assistance. Because each condition was evaluated twice in the double-reversed order, the average was calculated across both measurements of the condition. The cost of quiet standing was subtracted from the measured cost of all the other conditions to calculate the cost of walking. Users fasted for at least two hours before optimizations and at least four hours before validations to minimize the possible thermal effects of food on metabolic cost measurements. Two-tailed paired t-tests were used to evaluate if the metabolic cost of walking with exoskeleton assistance was significantly different from walking in the control conditions.

For the two-joint optimizations, the participant wore a cloth mask underneath the metabolics mask to comply with COVID-19 safety protocols. This mask did affect the metabolics measurements, by seeming to create a downward offset in the measured metabolic cost (Sup. Note 17). While this disrupted the accuracy of the absolute measurements, we expected the percent reductions in metabolic cost to be accurate, because we are comparing between conditions. We excluded these affected measures when calculating the average absolute measurements reported for standing quietly, walking with no exoskeleton, and walking with no torque.

#### Torques and Kinematics

Applied torques were measured using load cells and strain gauges on the exoskeleton. Exoskeleton joint angles were recorded to estimate user kinematics, meaning that we could not calculate kinematics for walking without the exoskeleton. Stride frequency was calculated using vertical ground reaction forces measured by the instrumented treadmill (Bertec). Measurements were averaged over the last three minutes of the condition for walking with no exoskeleton, and averaged over the last five minutes for walking with no torque and walking with each assistance condition. Measurements were averaged across both legs. For conditions that were evaluated twice (walking with no exoskeleton and walking with no torque), results were averaged across the two conditions.

Biological torques reported for reference (Fig. 3) were from a separate study. Reference walking data is from 3 healthy male subjects (38), different from the participants included in this study. Walking data was collected from 3 gait cycles of motion capture data during treadmill walking at 1.25 m/s including marker trajectories, ground reaction forces, and EMG measurements (38). Biological joint torques during walking were calculated using the Inverse Dynamics tool in OpenSim (55), as described in (25).

#### Muscle Activity

Muscle activity was measured using surface EMG (Delsys Trigno). We applied a 3rd order bandpass filter of 40 to 450 Hz, rectified, then applied a 3rd order low pass filter of 10 Hz (56). Muscle activity was averaged for a stride over the last five minutes of the device conditions and over the last three minutes for the no device conditions. We subtracted the baseline noise offset then normalized to the maximum of the no torque condition profile. The sensor locations are similar to the protocol of previous gait analysis experiments (21) with adjustments to avoid interfering with the device structure and straps.

## ACKNOWLEDGEMENTS

We would like to thank K. Gregorczyk, G. Kanagaki, M. O’Donovan and the NSRDEC for their input on experimental design, N. Bianco for assistance in controller development and reference biological torques, and all of the Stanford Biomechatronics Lab for their feedback and support. We would also like to thank the staff and administrators who were able to reopen the lab during the COVID-19 pandemic. This work was supported by the U.S. Army Natick Soldier Research, Development and Engineering Center (Grant number W911QY18C0140), by the National Science Foundation Graduate Research Fellowship Program (Grant number DGE-1656518), and by the Stanford Vice Provost for Undergraduate Education STEM Fellowship.

## Author Contributions

P. F. and G.B. designed and constructed the exoskeleton and developed the controllers. P.F. developed the optimization strategy and the experimental protocol, conducted experiments, and drafted and edited the manuscript. G.B., R.M., and R.R. also conducted experiments and edited the manuscript. S.C. conceived and managed the project, provided design, controls and testing support, and edited the manuscript.

---

**Supplementary Note 1: User Training**

**Supplementary Note 2: Metabolics - single-joint and Whole-leg**

**Supplementary Note 3: Metabolics - two-joint**

**Supplementary Note 4: Applied torques for each participant**

**Supplementary Note 5: Optimized parameters and parameter ranges**

**Supplementary Note 6: Applied exoskeleton power**

**Supplementary Note 7: Kinematics for each participant**

**Supplementary Note 8: Stride frequency for each participant**

**Supplementary Note 9: Muscle activity for each participant**

**Supplementary Note 10: Mobile device mass estimate**

**Supplementary Note 11: Experiment log - generations and reductions per day**

**Supplementary Note 12: Sample size and power analysis**

**Supplementary Note 13: Torque-tracking**

**Supplementary Note 14: Pilot tests**

**Supplementary Note 15: Detailed optimization methods**

**Supplementary Note 16: Tested parameter values during optimization**

**Supplementary Note 17: COVID mask effect on metabolics**

